# Mutation of *kvrA* causes OmpK35/36 porin downregulation and reduced meropenem/vaborbactam susceptibility in KPC-producing *Klebsiella pneumoniae*

**DOI:** 10.1101/845925

**Authors:** Punyawee Dulyayangkul, Wan Ahmad Kamil Wan Nur Ismah, Edward J. A. Douglas, Matthew B. Avison

## Abstract

Meropenem/vaborbactam resistance in *Klebsiella pneumoniae* is associated with loss of function mutations in the OmpK36 porin. Here we identify two previously unknown loss of function mutations that confer cefuroxime resistance in *K. pneumoniae*. The proteins lost were NlpD and KvrA; the latter is a transcriptional repressor that controls capsule production. We demonstrate that KvrA loss reduces OmpK35 and OmpK36 porin production, which confers reduced susceptibility to meropenem/vaborbactam in a KPC-3 producing *K. pneumoniae* clinical isolate.

## Text

Carbapenems are often reserved as a last resort for treatment of severe infections caused by multi-drug resistant Gram-negative bacteria. A rise in the prevalence of cephalosporin resistance, particularly due to the spread of mobile cephalosporinase genes in the Enterobacteriaceae has resulted in the increased use of carbapenems worldwide. This has driven the emergence of carbapenem resistant Enterobacteriaceae (CRE) which are classed as one of the greatest threats to human health according to the World Health Organisation. In *Klebsiella pneumoniae*, carbapenem resistance is mainly mediated by the production of a carbapenemase. In some parts of the world the most prevalent is the class A carbapenemase KPC; in others, most common are the class B metallo-β-lactamases e.g. NDM; in others, most class D carbapenemases e.g. OXA-48 like enzymes predominate (1–3). CRE can also emerge due to mutations that reduce envelope permeability, for example those that result in porin deficiency, and particularly when these mutations occur in isolates producing CTX-M or AmpC-type cephalosporinases (4). Indeed, most carbapenem resistant *K. pneumoniae* clinical isolates have multiple resistance mechanisms, including permeability defects (5–8).

As part of our work aiming to identify novel mechanisms for reduced cephalosporin and/or carbapenem permeability in *K. pneumoniae*, the *oqxR* and *ramR* double mutant, FQ3, derived from *K. pneumoniae* Ecl8 as previously described (9) was used as a parent strain to select cephalosporin resistant mutants. FQ3 overproduces two efflux pumps; AcrAB and OqxAB, resulting in resistance to fluoroquinolones, chloramphenicol and minocycline (9). One hundred microlitre aliquots of overnight cultures grown in Nutrient Broth (NB) were spread onto Mueller Hinton Agar containing 16 µg.mL^−1^ cefuroxime, which were then incubated for 24 h. One representative mutant derivative was named “FQ3 M1” and was shown by disc susceptibility testing, performed and interpreted according to CLSI methodology (10, 11) to be resistant to cefuroxime, cefoxitin and cephalexin (Table 1).

**Table 1:** Inhibition zone diameters (mm) for antimicrobials against *K. pneumoniae* FQ3 and cefuroxime resistant mutant FQ3 M1.

| Antimicrobial | Parental strain, FQ3 | FQ3 M1 |
| --- | --- | --- |
| Meropenem | 29 | 26 |
| Ertapenem | 28 | 23 |
| Imipenem | 28 | 25 |
| Doripenem | 27 | 26 |
| Aztreonam | 32 | 31 |
| Cefepime | 30 | 26 |
| Ceftazidime | 28 | 26 |
| Cefotaxime | 28 | 25 |
| Ceftriaxone | 30 | 27 |
| Ceftizoxime | 33 | 30 |
| Cefoperazone | 28 | 24 |
| Cefotetan | 25 | 21 |
| Cefoxitin | 19 | 10 |
| Cefuroxime | 19 | 13 |
| Cephalexin | 18 | 10 |
| Ciprofloxacin | 11 | 11 |
| Norfloxacin | 8 | 6 |
| Ofloxacin | 8 | 6 |
| Levofloxacin | 11 | 9 |
| Chloramphenicol | 8 | 6 |
| Minocycline | 13 | 12 |
| Amikacin | 26 | 25 |
| Gentamicin | 25 | 23 |

Values reported are the means of three repetitions rounded to the nearest integer for the diameter of the growth inhibition zone across each antimicrobial disc (mm). Shading indicates resistance according to susceptibility breakpoints set by the CLSI (11).

Changes in envelope protein abundance in mutant FQ3 M1 relative to FQ3 were quantified using LC-MS/MS proteomics as previously described (9) to identify potential reasons for cephalosporin resistance. There were 15 proteins significantly altered (Table 2). Amongst this group of proteins with uncertain or metabolic functions, were the important antimicrobial entry porins OmpK35 and OmpK36, which were both downregulated in FQ3 M1 by almost 3-fold.

**Table 2:** Significant changes in envelope protein abundance seen in *K. pneumoniae* mutant FQ3 M1 vs parent strain FQ3.

| Accession | Description | Abundance<br>FQ3 (1) | Abundance<br>FQ3 (2) | Abundance<br>FQ3 (3) | Abundance<br>FQ3 M1 (1) | Abundance<br>FQ3 M1 (2) | Abundance<br>FQ3 M1 (3) | T-test | Fold<br>change |
| --- | --- | --- | --- | --- | --- | --- | --- | --- | --- |
| A6T518 | Phosphoheptose isomerase, GmhA | - | - | - | 1.18E+08 | 2.05E+07 | 2.02E+07 | <0.0005 | >20 |
| A6T720 | Aspartate aminotransferase, AspC | - | - | - | 2.11E+07 | 1.92E+07 | 2.27E+07 | <0.0005 | >20 |
| A6TBN4 | ATP-binding component of methyl-galactoside transporter, MglA | - | - | - | 2.64E+07 | 2.11E+07 | 3.40E+07 | <0.0005 | >20 |
| A6TF45 | sn-glycerol-3-phosphate dehydrogenase, GlpD | 3.10E+07 | 1.21E+07 | 3.24E+07 | 2.37E+08 | 1.24E+09 | 1.72E+09 | 0.004 | 42.43 |
| A6T5Q5 | Putative outer membrane protein | 4.62E+08 | 3.06E+08 | 4.48E+08 | 1.61E+09 | 1.85E+09 | 1.51E+09 | <0.0005 | 4.09 |
| A6TBM3 | Putative channel/filament proteins, YohG | 2.93E+08 | 2.57E+08 | 4.77E+08 | 9.04E+08 | 1.04E+09 | 1.31E+09 | <0.0005 | 3.16 |
| A6T4Y7 | Acetyl-coenzyme A carboxylase carboxyl transferase, AccA | 1.24E+08 | 8.31E+08 | 4.06E+08 | 1.15E+09 | 7.38E+08 | 9.36E+08 | 0.005 | 2.07 |
| A6T8N5 | Putative uncharacterized protein, YdgH | 6.65E+08 | 5.22E+08 | 8.53E+08 | 3.69E+08 | 3.61E+08 | 3.44E+08 | 0.001 | 0.53 |
| A6T721 | OmpF porin (OmpK35) | 1.19E+10 | 7.32E+09 | 7.82E+09 | 2.89E+09 | 3.97E+09 | 3.01E+09 | 0.001 | 0.36 |
| A6TBT2 | OmpC porin (OmpK36) | 3.49E+10 | 3.13E+10 | 3.41E+10 | 1.12E+10 | 1.31E+10 | 1.00E+10 | <0.0005 | 0.34 |
| A6TD34 | Lipoprotein, NlpD | 9.15E+08 | 7.51E+08 | 1.14E+09 | 2.64E+08 | 1.68E+08 | 3.13E+08 | <0.0005 | 0.27 |
| A6T4Y4 | Lipid-A-disaccharide synthase, LpxB | 6.06E+07 | 5.19E+07 | 1.02E+08 | - | - | - | <0.0005 | <0.005 |
| A6T9Z3 | Putative multidrug resistance protein, YdhJ | 7.23E+07 | 6.50E+07 | 9.18E+07 | - | - | - | <0.0005 | <0.005 |
| A6TBM5 | Putative cellobiose-specific PTS permease | 2.83E+07 | 1.66E+07 | 3.57E+07 | - | - | - | <0.0005 | <0.005 |
| A6TEA5 | Putative glycerol dehydrogenase, DhaD | 5.00E+07 | 2.28E+08 | 2.51E+08 | - | - | - | <0.0005 | <0.005 |

Strains were grown in nutrient broth and raw abundance data are provided for three biological replicates of parent (FQ3) and mutant (FQ3 M1).
Analysis was as described in (9) and proteins listed are those with significantly different abundance in FQ3 M1 versus FQ3.

We next used WGS to try and explain the downregulation of OmpK35 and OmpK36 in mutant FQ3 M1 relative to FQ3. WGS was performed by MicrobesNG (Birmingham, UK) using a HiSeq 2500 instrument (Illumina, San Diego, CA, USA). Reads were trimmed using Trimmomatic (12) and assembled into contigs using SPAdes 3.10.1 (http://cab.spbu.ru/software/spades/). Assembled contigs were mapped to the *K. pneumoniae* Ecl8 reference genome (GenBank accession number GCF_000315385.1) by using progressive Mauve alignment software (13).

No mutations were detected in *ompK36* and *ompK35* or adjacent sequences, or in known regulators of porin production, e.g. OmpR/EnvZ (14). In fact, FQ3 M1 has three separate mutations relative to FQ3: a frameshift mutation (causing Asn278FS) in *nlpD*, a frameshift (causing Asp395FS) in *dhaR*, and a 1,159 bp deletion spanning *kvrA* and the adjacent genes, *ydhI* and *ydhJ*. In *Escherichia coli*, NlpD is involved in peptidoglycan remodelling during cell division (15–17). DhaR is a transcriptional activator responsible for controlling the production of the metabolic enzymes glycerol dehydrogenase and dihydroxyacetone kinase (18,19). KvrA is a MarR-type transcriptional repressor with a key role in *K. pneumoniae* capsulation (20–22).

In order to deconvolute the possible roles of the three different loss of function mutations, we separately insertionally inactivated *nlpD*, *dhaR* and *kvrA*, plus *ompK36* as controls in FQ3. Mutants were constructed using the pKNOCK suicide plasmid (23). DNA fragments of the genes to be inactivated were amplified with Phusion High-Fidelity DNA Polymerase (NEB, UK) from *K. pneumoniae* Ecl8 genomic DNA by using primers *ompK36* KO FW (5′-CGTTCAGGCGAACAACACTG-3′) and *ompK36* KO RV (5′-AAGTTCAGGCCGTCAACCAG-3′); *kvrA* KO FW (5’-ATCTGGCACGTTTAGTTCGC-3’) and *kvrA* KO RV (5’-CCCTTTCTCCTCCAGCTGAT-3’); *dhaR* KO FW (5’-CAATCAGATGTACGGCCTGC-3’) and *dhaR* KO RV (5’-GACTTCGACGTGATTCAGGC-3’); *nlpD* KO FW (5’-ACGATTTCCGCGACCTGGCG-3’) and *nlpD* KO RV (5’-CAACATCTTGGTAGCACTCT-3’). Each PCR product was separately ligated into the pKNOCK-GM at the SmaI site and each recombinant plasmid was then transferred into FQ3 cells by conjugation from *E. coli* BW20767. Mutants were selected using gentamicin (5 µg.mL^−1^) and the mutations were confirmed by PCR using primers *ompK36* full length FW (5’-GAGGCATCCGGTTGAAATAG-3’) and *ompK36* full length RV (5’-ATTAATCGAGGCTCCTCTTAC-3’); *kvrA* full length FW (5’-ACTTAGCAAGCTAATTATAAGGAGATGA-3’) and *kvrA* full length RV (5’-GCCGCAAAGAATTAATCTTTA-3’); *dhaR* full length FW (5’-CAGCCCGATGGACGAGATT-3’) and *dhaR* full length RV (5’-TATTGGGCTCAGCGCGTCC-3’); *nlpD* full length FW (5’-GTCGGCGAAGAGCATCAGT-3’) and *nlpD* full length RV (5’-CACCTTCCACGGCACATCA-3’).

Inactivating *dhaR* in FQ3 had no effect on cephalosporin MIC, determined using CLSI methodology (24) but inactivating *nlpD* or *kvrA*, raised cefoxitin and cefuroxime MIC, though in both cases the MIC was one doubling lower than against FQ3 M1 (Table 3). We next complemented *kvrA* in FQ3 M1. To do this, *kvrA* DNA was amplified from *K. pneumoniae* Ecl8 genomic DNA by using primers (introduced restriction sites underlined): *kvrA* full length *Bam*HI FW (5’-AAA<u>GGATCC</u>CGGCAATCCGGATGTGTTAAGAC-3’) and *kvrA* full length SalI RV (5’-AAA<u>GTCGAC</u>GGAGGGTGAAAAAAGGCCCGGATTA-3’). The PCR product was digested and inserted to pUBYT (25) cut with BamHI and SalI to generate pUBYT::*kvrA*. The recombinant plasmid was then transferred into FQ3 M1 cells by electroporation. The transformants were selected using kanamycin (50 µg.mL^−1^) and the presence of plasmids was confirmed by PCR using primers pUBYT check FW (5’-GCAAGAAGGTGATGAATCTACA-3’) and pUBYT check RV (5’-GTGGCAGCAGCCAACTCA-3’). Complementation of FQ3 M1 with pUBYT::*kvrA* showed that the MICs of cefoxitin and cefuroxime reduced, but again not to a value as low as against FQ3. This confirmed the result of the gene inactivation experiment that *kvrA* loss alone is not the sole determinant of the cephalosporin resistant phenotype expressed by FQ3 M1 (Table 3).

**Table 3.** MICs (µg.mL^−1^) of cephalosporins against *K. pneumoniae* Ecl8 FQ3 and mutant derivatives.

|  | Cefuroxime MIC | Cefoxitin MIC |
| --- | --- | --- |
| FQ3 | 8 | 8 |
| FQ3 $\Delta dhaR$ | 8 | 8 |
| FQ3 $\Delta ompK35$ | 8 | 8 |
| FQ3 $\Delta kvrA$ | 16 | 32 |
| FQ3 $\Delta nlpD$ | 32 | 32 |
| FQ3 $\Delta ompK36$ | 32 | 32 |
| FQ3 M1 | 64 | 64 |
| FQ3 M1(pUBYT) | 32 | 64 |
| FQ3 M1(pUBYT::*kvrA*) | 16 | 16 |

The CLSI susceptible and resistance breakpoints (11) for cefuroxime and cefoxitin are $\leq 8$ and $\geq 32 \mu\text{g.mL}^{-1}$ .

Given that inactivation of either *kvrA* or *nlpD* in FQ3 reduced cephalosporin susceptibility, we performed LC-MS/MS envelope proteomics for these mutants versus FQ3 as above, which revealed that mutation in *kvrA* caused a reduction in OmpF (OmpK35) and OmpC (OmpK36) porin levels in FQ3, to the same extent as seen in FQ3 M1. Mutation in *nlpD*, despite altering cephalosporin MIC, did not (Fig. 1 A, B). Because FQ3 is derived from Ecl8, a laboratory strain, and because FQ3 carries mutations that increase efflux pump production (9), we wanted to test the impact of *kvrA* inactivation in a wild-type clinical isolate. To do this, we insertionally inactivated *kvrA* (as above) in the susceptible *K. pneumoniae* clinical isolate S17, which has wild-type envelope permeability (26) and showed by LC-MS/MS that OmpF and OmpC levels fell in this mutant relative to S17, as seen in FQ3. Porin abundance did not change upon *nlpD* inactivation in S17, also as seen in FQ3 (Fig. 1 C, D).

**Figure 1.**
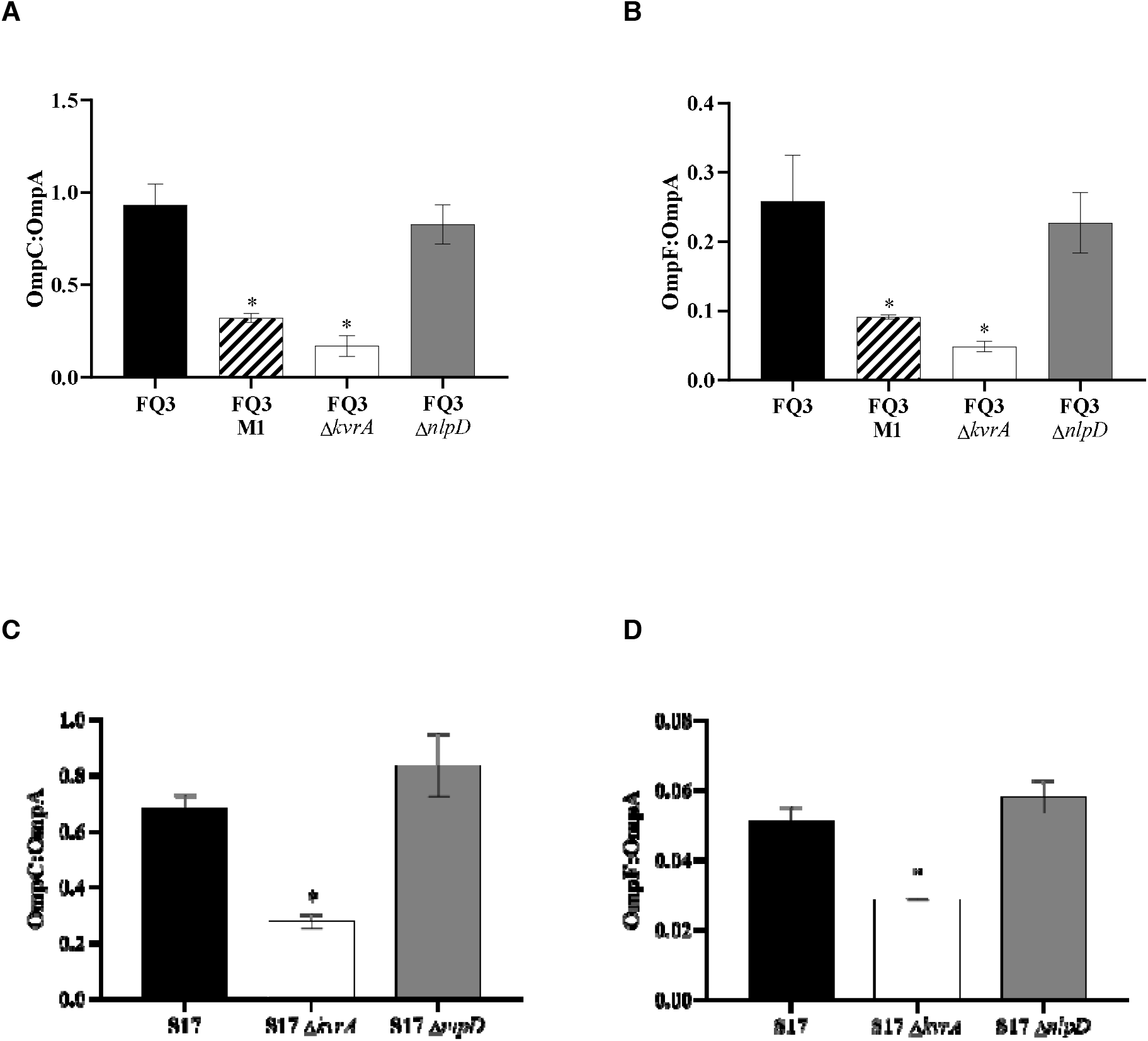
Normalised abundance of OmpK35 and OmpK36 porins in *K. pneumoniae kvrA* and *nlpD* mutants. OmpC (OmpK36) and OmpF (OmpK35) abundance was measured using LC-MS/MS and normalised to the abundance of OmpA, to control for protein loading. Data presents mean ± standard error of the mean, *n*=3. Asterisk (*) indicates statistically significant differences from the parental strains by t-test (*p*<0.05). (**A**) OmpC to OmpA protein ratio and (**B**) OmpF to OmpA protein ratio in FQ3 and its mutant derivatives. (**C**) OmpC to OmpA protein ratio and (**D**) OmpF to OmpA protein ratio in clinical isolate S17 and its mutant derivatives.

β-Lactamase inhibitors such as clavulanic acid, tazobactam and sulbactam have been successful in overcoming resistance to penicillin derivatives, e.g. amoxicillin, piperacillin and ticarcillin in Enterobacteriaceae. However, inhibitor/penicillin combinations are not clinically useful against KPC, CTX-M, OXA-48-like or AmpC producing Enterobacteriaceae isolates, or those producing metallo-β-lactamases (27). This has led to the development of new β-lactam/β-lactamase inhibitor combinations, and one recently licenced for clinical use is meropenem/vaborbactam (28). Vaborbactam is a serine β-lactamases inhibitor based on a cyclic boronic acid pharmacophore. It has potent in vitro activity against class A β-lactamases, particularly KPC, and it can restore the activity of meropenem against KPC-producing Enterobacteriaceae (29–31).

Meropenem/vaborbactam resistance in KPC-producing *K. pneumoniae* has been shown to occur by loss of function mutation in *ompK36*, which encodes is the primary porin for meropenem entry (32,33). We therefore wanted to test the impact of OmpK36 porin reduction seen following inactivation of *kvrA* in *K. pneumoniae* S17 (Fig. 1C) on meropenem/vaborbactam susceptibility when the mutant produces KPC. To investigate this, we used pUBYT::*bla*_KPC-3_ (25) to transform *K. pneumoniae* S17, S17Δ*kvrA* and S17Δ*ompK36* (as a control). As expected, due to carriage of KPC, all transformants were resistant to meropenem. Addition of 8 µg.mL^−1^ vaborbactam (8 µg.mL^−1^) reduced the meropenem MIC against S17 well into the susceptible range. In contrast, meropenem/vaborbactam MICs against the *kvrA* and *ompK36* mutants were 1 µg.mL^−1^ (four doublings higher than against S17) and 32 µg.mL^−1^ respectively (Table 4). Hence, even in an otherwise wild-type KPC-producing clinical *K. pneumoniae* isolate (26), OmpK35/OmpK36 downregulation caused solely by *kvrA* mutation is enough to reduce meropenem/varborbactam susceptibility.

**Table 4.**
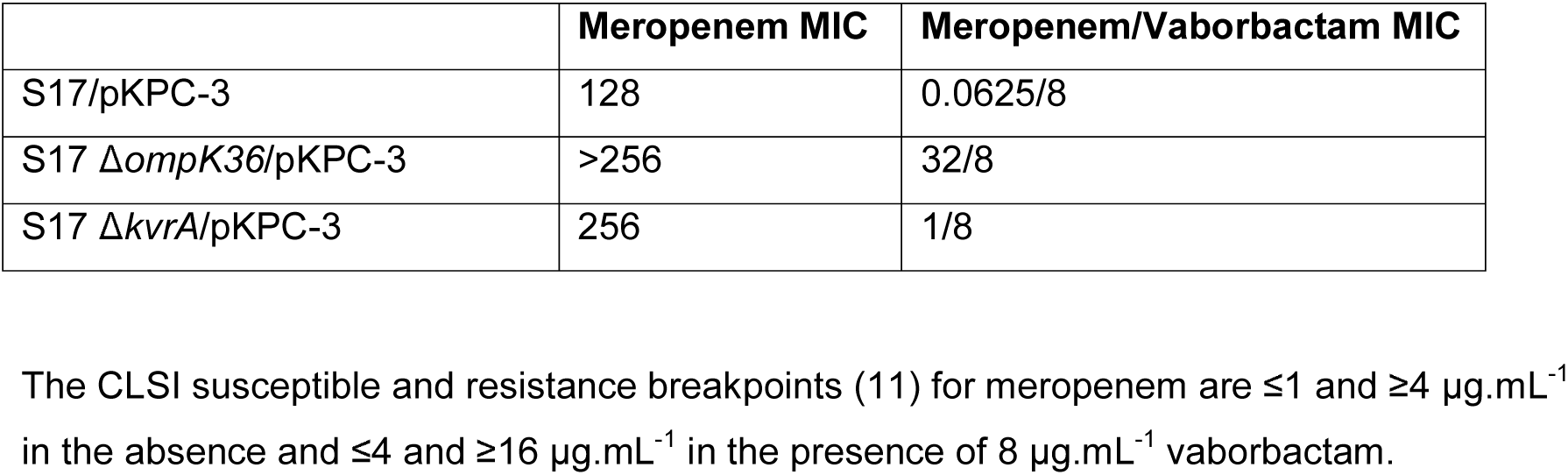
MICs (µg.mL^−1^) of meropenem against *K. pneumoniae* clinical isolate S17 and mutant derivatives measured with and without 8 µg.mL^−1^ of vaborbactam.

|  | **Meropenem MIC** | **Meropenem/Vaborbactam MIC** |
| --- | --- | --- |
| S17/pKPC-3 | 128 | 0.0625/8 |
| S17 $\Delta ompK36$/pKPC-3 | >256 | 32/8 |
| S17 $\Delta kvrA$/pKPC-3 | 256 | 1/8 |

The CLSI susceptible and resistance breakpoints (11) for meropenem are $\leq 1$ and $\geq 4$ $\mu\text{g.mL}^{-1}$ in the absence and $\leq 4$ and $\geq 16$ $\mu\text{g.mL}^{-1}$ in the presence of 8 $\mu\text{g.mL}^{-1}$ vaborbactam.

In conclusion, we report here two loss of function mutations in genes previously not known to affect antimicrobial susceptibility in *K. pneumoniae*. NlpD (“new lipoprotein D”) is conserved across Gram-negative bacteria, with essential roles in virulence, e.g. in *Yersinia pestis* (34). In *E. coli* it is recruited to the division site where it targets the activity of the peptidoglycan amidase AmiC, to which it binds. Loss of NlpD in *E. coli* is known to delay the onset of cell lysis after treatment with ampicillin because peptidoglycan breakdown by AmiC is less targeted to the division site (35,36). This provides a clear rationale for why disruption of *nlpD* reduces β-lactam susceptibility in *K. pneumoniae* (Table 1), but its role as a mediator of cefuroxime resistance has not previously been suspected.

We also report that inactivation of *kvrA* causes cefuroxime resistance in *K. pneumoniae*, but more importantly, it causes reduced susceptibility meropenem/vaborbactam, even in an otherwise wild-type *K. pneumoniae* clinical isolate, transformed to express *bla*_KPC-3_ from its native promoter in a low copy number vector (25). We show that cefuroxime resistance and reduced meropenem/vaborbactam susceptibility in a *kvrA* mutant is associated with OmpK35/OmpK36 downregulation. This is reminiscent of OmpR mutations in *E. coli*, which reduce OmpC/OmpF production and can also affect antimicrobial susceptibility (37).

KvrA is a MarR-family transcriptional repressor. Importantly, we found that YdhJ is upregulated 45-fold (*p*=0.002, *n*=3) and >100-fold (*p*<0.0001, *n*=3) according to proteomics following disruption of *kvrA* in *K. pneumoniae* S17 and FQ3, respectively. YdhJ is encoded within a putative efflux pump operon adjacent to *kvrA* on the chromosome. Expression of the homologue of this *ydhIJK* operon in *E. coli* is directly repressed by SlyA (38), which is encoded upstream of *ydhIJK*, in an almost identical arrangement as *kvrA* and *ydhIJK* in *K. pneumoniae*. Therefore, the *ydhIJK* operon is likely to be the direct repressive target of KvrA in this species, with its wider activatory effects being indirect. Similar characterised MarR-family repressors such as SlyA also tend to have local direct repressive effects, but cause activation of gene expression at some promoters indirectly by blocking the repressive activity of H-NS at those promoters (39). Since H-NS is known to have a repressive effect on porin production in *E. coli* (40) it may well be that increased repressive activity of H-NS in the absence of KvrA is the explanation for OmpK35/OmpK36 downregulation seen in *K. pneumoniae*.

Disruption of *kvrA* also causes downregulation of key capsule biosynthesis genes, e.g. *galF*, *manC* and *wzi* in some *K. pneumoniae* strains (20). We found using proteomics that Wzi was downregulated a marginal 0.82-fold (*p*=0.03, *n*=3) upon disruption of *kvrA* in isolate S17 but not the GalF or ManC and none of these proteins were downregulated in FQ3Δ*kvrA* versus FQ3. However, it is known that *kvrA* loss does not reduce capsule production is all strains, and nor does it attenuate the virulence of all strains in a mouse infection model (20). Among 455 isolates of *K. pneumoniae* from NCBI database, we found 9 isolates (Genbank accession numbers CP037927, LR134162, CP032175, CP018056, CP003200, LR588409, CP044039, CP043670, CP043669) where *kvrA* was mutated in a way predicted to result in a truncated and presumably non-functional KvrA. One key example is the KPC-2 and CTX-M-14 producing carbapenem resistant human sputum isolate HS11286 (41, 42). This confirms that *kvrA* mutants are present in human clinical samples.

## Acknowledgments

This work was funded by grant MR/S004769/1 to M.B.A. from the Antimicrobial Resistance Cross Council Initiative supported by the seven United Kingdom research councils and the National Institute for Health Research. Genome sequencing was provided by MicrobesNG (http://www.microbesng.uk/), which is supported by the BBSRC (grant number BB/L024209/1). W.A.K.W.N.I. was funded by a postgraduate scholarship from the Malaysian Ministry of Education. We are grateful to Dr Kate Heesom, School of Biochemistry, University of Bristol for performing the proteomics analysis and Melissa Rose Bennett, School of Cellular & Molecular Medicine, University of Bristol for assistance in selecting FQ3 M1.

**We declare no conflicts of interest.**

